# Visualizing and exploring patterns of large mutational events with SigProfilerMatrixGenerator

**DOI:** 10.1101/2023.02.03.527015

**Authors:** Azhar Khandekar, Raviteja Vangara, Mark Barnes, Marcos Díaz-Gay, Ammal Abbasi, Erik N. Bergstrom, Christopher D. Steele, Nischalan Pillay, Ludmil B. Alexandrov

## Abstract

**Background:** All cancers harbor somatic mutations in their genomes. In principle, mutations affecting between one and fifty base pairs are generally classified as small mutational events. Conversely, large mutational events affect more than fifty base pairs, and, in most cases, they encompass copy-number and structural variants affecting many thousands of base pairs. Prior studies have demonstrated that examining patterns of somatic mutations can be leveraged to provide both biological and clinical insights, thus, resulting in an extensive repertoire of tools for evaluating small mutational events. Recently, classification schemas for examining large-scale mutational events have emerged and shown their utility across the spectrum of human cancers. However, there has been no standard bioinformatics tool that allows visualizing and exploring these large-scale mutational events

**Results:** Here, we present a new version of SigProfilerMatrixGenerator that now delivers integrated capabilities for examining large mutational events. The tool provides support for examining copy-number variants and structural variants under two previously developed classification schemas and it supports data from numerous algorithms and data modalities. SigProfilerMatrixGenerator is written in Python with an R wrapper package provided for users that prefer working in an R environment.

**Conclusions:** The new version of SigProfilerMatrixGenerator provides the first standardized bioinformatics tool for optimized exploration and visualization of two previously developed classification schemas for copy number and structural variants. The tool is freely available at https://github.com/AlexandrovLab/SigProfilerMatrixGenerator with an extensive documentation at https://osf.io/s93d5/wiki/home/.

## BACKGROUND

Large-scale cancer genomics projects have comprehensively surveyed the molecular landscapes of most types of human cancer [1, 2]. These studies have provided a compendium of somatic mutations for each examined cancer genome and revealed both the mutations driving cancer development and the processes generating most somatic mutations within each cancer [1–3]. One commonly performed type of genomics analysis is the examination of mutational patterns within a set of cancer genomes and the extraction of mutational signatures that have given rise to these patterns [3, 4]. Historically, mutational patterns have been predominately examined in the context of small mutational events, which include single base substitutions (SBS), doublet base substitutions (DBS), and small insertions and deletions (IDs) [3, 5]. Recent studies have also started exploring the patterns of large mutational events, including ones due to copy-number alterations and/or structural variations [6, 7]. Previously, we developed a computational tool, termed, SigProfilerMatrixGenerator, designed exclusively for examining the mutational patterns of all types of small mutational events [8]. Here, we present a new version of SigProfilerMatrixGenerator that now provides the capabilities for optimized exploration and visualization of large mutational events.

Large mutational events, generally defined as genomic alterations greater than 50 base pairs, are an important class of somatic aberrations in human cancer [6]. In principle, there are two commonly examined and closely interrelated types of large mutational events: *(i)* a structural variation (SV, also known as a genomic rearrangement), where a large-scale genomic segment gets altered; and *(ii)* a copy number variation (CNV), where the number of DNA copies of a genomic segment gets modified. Not all structural variations are related to CNVs, as SVs do not necessarily alter the copy number of a genomic segment; examples include copy neutral events such as inversions and reciprocal translocations. Similarly, not all changes in copy number require prior SVs, as is the case of chromosomal duplications and whole-genome doubling. Importantly, SVs and CNVs also differ in the types of genomics approaches that can detect them. In most cases, comprehensive detection of SVs requires whole-genome sequencing (WGS) data as it relies on either read alignment [9] or genome assembly methods [10]. In contrast, in addition to WGS data, CNVs can be detected from whole-exome sequencing, RNA-sequencing, single-cell sequencings approaches, and genotyping microarrays [11–13].

Deciphering mutational signatures from catalogues of somatic mutations, a process known as *de novo* signature extraction, relies on a biologically meaningful classification of mutational events [5]. We previously created the mathematical concept of mutational signatures and provided a set of tools for deciphering signatures of small mutational [4, 8]. Mutational patterns of SBSs, DBSs, IDs, have been extensively explored with more than 100 distinct mutational signatures published in the literature [3, 14]. These signatures reflect the activities of endogenous and/or exogenous mutational processes with an approximately half of all signatures being, at least putatively, linked with a proposed etiology [15–18]. Recently, mutational signature analyses of larger copy number alterations and structural alterations have emerged [6, 7, 19, 20]. A crucial first step in extracting mutational signatures is the derivation of features according to a predefined schema for mutational classification. This step involves transforming the mutational catalogues of a set of cancer genomes into a matrix, which is then amenable to subsequent matrix decomposition techniques [8]. Here, we present a computational package for classification of large-scale alterations and the generation of mutational matrices for signature decomposition.

Two separate classification schemas are implemented: one for copy number variations and one for structural variations. Both schemas were previously developed and applied to large cohorts of cancer samples [7, 19, 21]. To the best of our knowledge, there is currently no tool that allows matrix generation and visualization of SVs and CNVs classified under these schemas. SigProfilerMatrixGenerator’s capabilities for analyzing SVs and CNVs are implemented in Python and the tool allows using multiple input formats, including segmentation and browser extensible data paired-end (BEDPE) files generated by commonly used algorithms for detecting copy number variations and structural variations, respectively. Additionally, SigProfilerMatrixGenerator provides a comprehensive visualization of mutational patterns of large mutational events and an R wrapper package for users that prefer working within the R environment.

## IMPLEMENTATION

### Classification of Copy Number Variations

The schema for classifying copy number variations is based on Steele *et al*. [7] and it utilizes allele-specific copy number, which quantifies the number of segments for each allele at each variant loci rather than the total number of chromosome copies. In this schema, the copy-number profile of a sample can be represented by a mutational vector with 48 dimensions. Specifically, copy number segments are categorized into three heterozygosity states: heterozygous segments with total copy number (TCN) of A>0, B>0 (numbers reflect the counts for major allele *A* and minor allele *B*; **Figure 1*a***), segments with loss of heterozygosity (LOH) with total copy number of A>0, B=0 (**Figure 1*b***), and segments with homozygous deletions and TCN of A=0, B=0 (**Figure 1*c***). Segments are further subclassified into 5 categories based on total copy number, which reflects the sum of the copies on the major allele *A* and the copies on the minor allele *B*: TCN=0, TCN=1, TCN=2, TCN=3 or 4, TCN=5 to 8, and TCN>=9. Each of these total copy number states accounts for the phenomenon of whole-genome duplication, for example a diploid (TCN=2) state transitioning to a doubled state (TCN=4), and a subsequent doubling of this state to TCN=8 is accounted for by the TCN=5-8 category (**Figure 1*a***). The categories for total copy number have been chosen for biological relevance (**Figure 1)**: TCN=0 reflects homozygous deletions, TCN=1 represents a genomic deletion resulting in an LOH, TCN=2 is equivalent to a diploid state including copy neutral LOH (a phenomenon whereby one of two homologous chromosomal regions is lost, but two identical copies of this region still remain; **Figure 1*b***), TCN=3 or 4 reflect a gained state of tri-to tetra-ploidy, TCN=5 to 8 represent a penta-to octo-ploidy state, and TCN>=9 represents high-level amplifications such as ones found in samples containing extrachromosomal DNA (ecDNA) [22]. Each of the heterozygous and LOH total copy number categories are additionally subclassified into five additional categories based on the size of their segments: 0 – 100kb, 100kb – 1Mb, 1Mb – 10Mb, 10Mb – 40Mb, and >40Mb. Three size bins are used for the additional subcategorization of homozygous deletions: 0 – 100kb, 100kb – 1Mb, and >1Mb. The partitioning by segment sizes was chosen to ensure that a sufficient proportion of segments are classified within each category [7]. This classification allows summarizing copy number profiles using 48 distinct channels and can be represented using a vector with 48 components. For example, a sample harboring multiple focal amplifications, either contained on linear or extrachromosomal DNA, will have many events in the 9+ total copy number category and the first 3 size bins (0 – 100kb, 100kb – 1Mb, 1Mb – 10Mb; **Figure 2*a-b***). Conversely, a sample containing a large number of focal deletions or losses of entire chromosomes or chromosome arms will have numerous events in the LOH category, spanning all size bins (**Figure 2*c-d***). Another example will be a sample with a whole-genome doubling where copy number changes will primarily encompass segments with large genomic sizes (10Mb – 40Mb; 40Mb) and total copy number between 3 and 4 (**Figure *2e-f*).** Overall, this 48-channel classification schema can effectively summarize a diverse array of copy number states seen across tumor types [7], whether they contain broad or focal events that result in amplifications or deletions.

**Figure 1.**
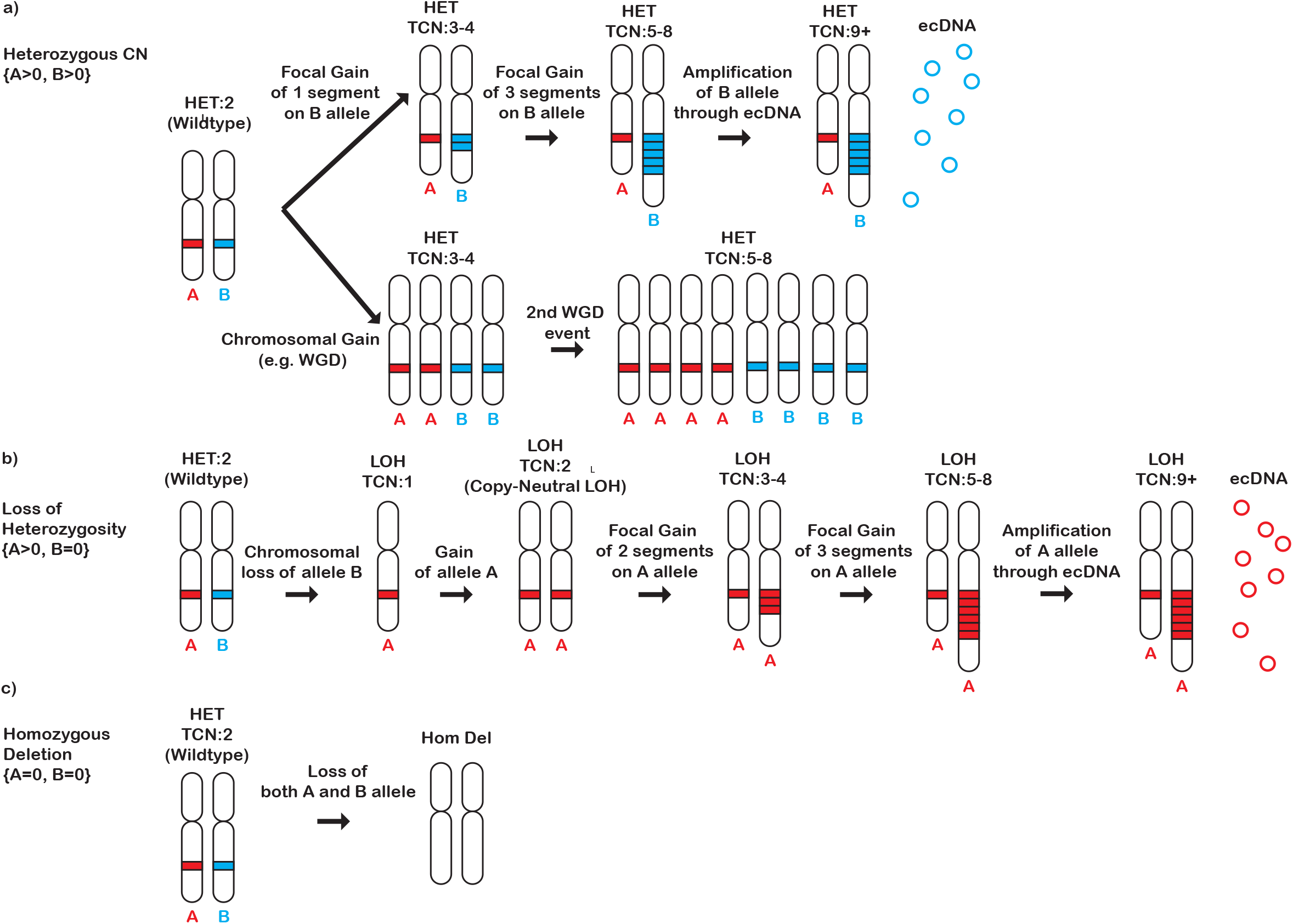
Description of the Copy Number Classification Schema. The copy number classification schema consists of 48 mutually exclusive channels, divided by heterozygosity status, segment size, and total copy number (TCN). ***a)*** In the heterozygous state, both alleles are retained and either one or both alleles can be amplified. This amplification can be focal (top panel) or it can encompass a chromosome or even the whole genome (bottom panel). The heterozygous category is further subdivided based on TCN (TCN=1, TCN=2, TCN=3 or 4, TCN=5 to 8, and TCN>=9). ***b)*** In a state of loss of heterozygosity (LOH), one of the alleles is lost. The remaining allele can then be duplicated (i.e., copy neutral LOH), and undergo more amplification resulting in higher total copy number states. The LOH category is further subdivided based on TCN (TCN=0, TCN=1, TCN=2, TCN=3 or 4, TCN=5 to 8, and TCN>=9). The heterozygous and LOH categories are further divided on the basis of the size of the segment: 0 – 100kb, 100kb – 1Mb, 1Mb – 10Mb, 10Mb – 40Mb, >40Mb. High-level LOH or heterozygous amplifications (e.g., TCN=5 to 8 or TCN>= 9) can be carried on extrachromosomal DNA (depicted as red circles) as well as on linear chromosomes. **c)** Homozygous deletions result in the loss of both alleles, and are divided on the basis of the size of the deleted segment: 0 – 100kb, 100kb – 1Mb, and >1Mb.

**Figure 2.**
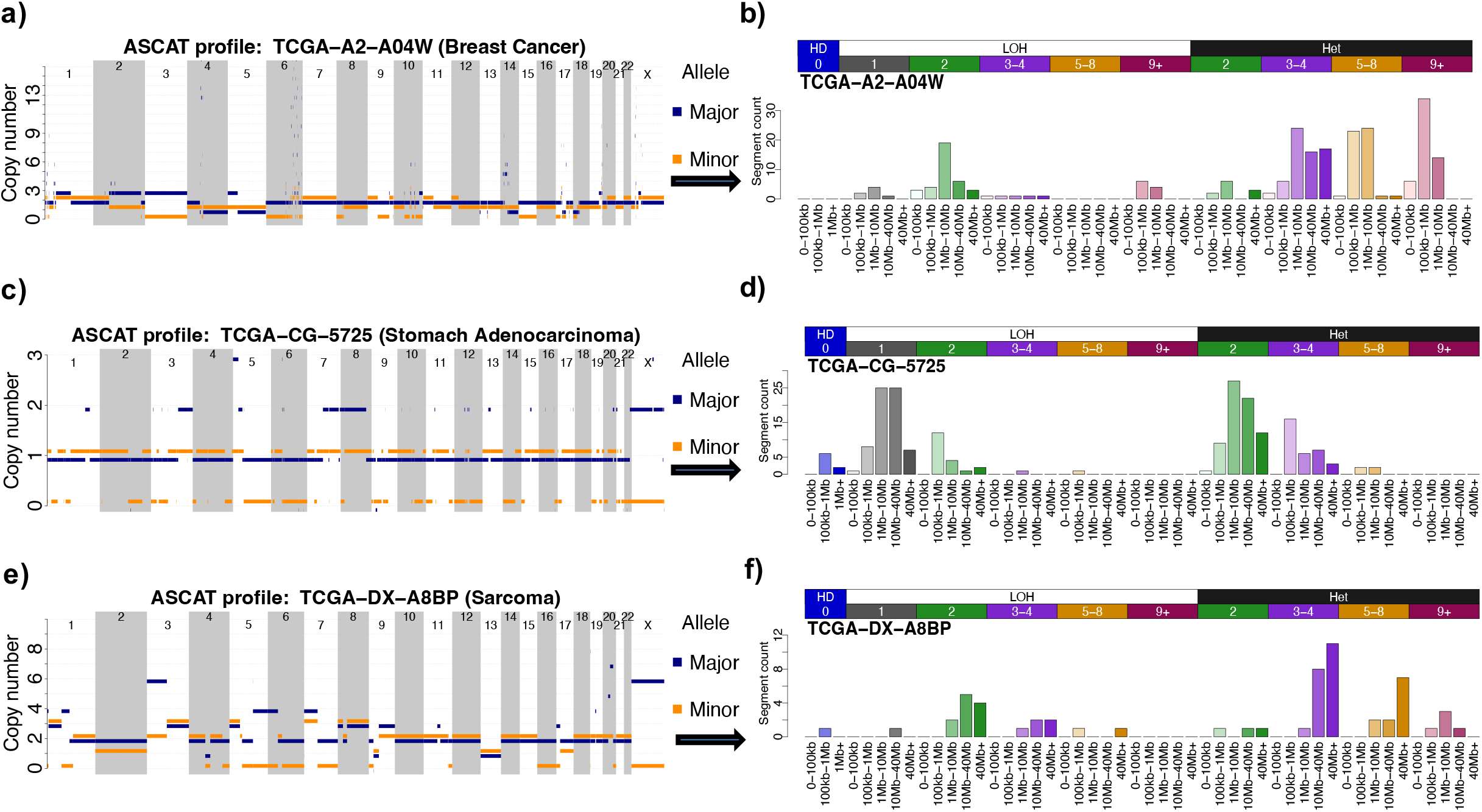
Converting Copy Number Segmentation Profiles into Copy Number Mutational Vectors. The CNV classification schema converts a sample’s segmentation profile (**a**, **c**, **e**) into a count vector of 48 mutually exclusive components (**b**, **d**, **f**). These components are based on segment size, heterozygosity status, and total copy number. A breast cancer sample with many highly amplified segments, possibly due to the presence of extrachromosomal DNA, is shown in (**a**, **b**). This sample’s count vector is characterized by peaks in the 5-8 and 9+ total copy number categories. A gastric cancer sample with extensive loss of heterozygosity is shown in (**c**, **d**). This sample’s count vector is characterized by peaks in the LOH category, specifically with a total copy number of 1 indicating a loss of an allele. A sarcoma sample with a whole-genome duplication event, characterized by peaks in the 3-4 total copy number category and the 40+ Mb size bin, is shown in (**e**, **f**).

### Input Data for Classifying Copy Number Variations

SigProfilerMatrixGenerator allows examining allele specific CNV data that, at a minimum, include the following information for each CNV segment: chromosome, start coordinate, end coordinate, and copy number of both the minor and major allele. Output files from the following tools for detecting CNVs are automatically supported: ASCAT [23], ABSOLUTE [24], Sequenza [25], FACETS [12], Battenberg [23], and PURPLE [26]. Additionally, custom segmentation files from other CNV detection tools can be used if these files contain the aforementioned information.

### Classification of Structural Variations

A classification schema consisting of 32 features, based on Nik-Zainal *et al*. [21], is used to construct a mutational vector with 32 dimensions for each sample. In principle, each structural variant consists of two breakpoints which are at single-base resolution, where a breakpoint is defined as a junction that indicates a structurally variable genomic segment greater than 50 base pairs [10]. Breakpoints are typically detected using three signals from aligned sequencing reads: depth of sequence coverage, discordant read-pairs, and split read-pairs [27–29]. Breakpoints can also be detected via genome assembly, where reads are assembled into contigs, the contigs are aligned to the reference genome, and these alignments are analyzed for structural variants [10]. The previously developed classification of structural variants considers the following canonical SVs: tandem duplications, deletions, inversions, and translocations (**Figure 3**). A tandem duplication refers to a segment of genomic material that has been duplicated and inserted on the same chromosome adjacent to the original segment (**Figure 3*a***). It should be noted that a tandem duplication is not necessarily the same as a copy-number amplification. For example, ecDNA copy-number amplifications are not tandem duplications as they are not inserted adjacent to the original chromosome segment. A somatic deletion is an event that has removed a set of existing base-pairs from a given location of a chromosome (**Figure 3*b***). An inversion is when a segment of the chromosome breaks off and reattaches at the same locus but in a reverse orientation (**Figure 3*c***). A translocation event occurs when a piece of one chromosome breaks off and some (or all) fragments from that piece re-attach to either another chromosome or to a different locus of the same chromosome (**Figure 3*d***). The classification schema bins all SVs, apart from translocations, according to the size of the event in base pairs: 0–10kb, 10kb–100kb, 100kb– 1Mb, 1Mb–10Mb, and >10Mb [21]. Translocations, which may involve more than one chromosome, are not binned by size because they can be either balanced (where there is no net loss of genetic material on the chromosomes involved and thus the size can be described by one number) or unbalanced (where there is a net loss or gain of genetic material on the chromosomes involved and thus the sizes of the segments cannot be described by just one number). Note that whether a translocation is balanced or unbalanced is not considered in this classification schema. The different types of SVs are then further divided into *clustered* and *non-clustered* events to account for the non-random distribution of these events along the genome. Clustered events are defined as events that occur closer to each other on a chromosome than purely expected by chance. These clusters often arise as a result of complex events, such as chromothripsis [30] or chromoplexy [31], generating many breakpoints in a single instantaneous event as opposed to the gradual accumulation of events over many cell cycles which results in more dispersed non-clustered events. Clusters of breakpoints can also form as a result of other mechanisms, including, for example, rearrangement hotspots in the genome [32]. Clustering of SVs is determined based on a previously developed algorithm that utilizes the Potts’ filter method [33]. This method segments a chromosome based on inter-mutational distance of SV breakpoints, and if the average distance in a particular segment is less than 10 times the average inter-mutational distance in the sample, all breakpoints in the segment are considered clustered. A minimum of 10 breakpoints must be present for a given segment to be considered clustered, otherwise all breakpoints in that segment are considered non-clustered.

**Figure 3.**
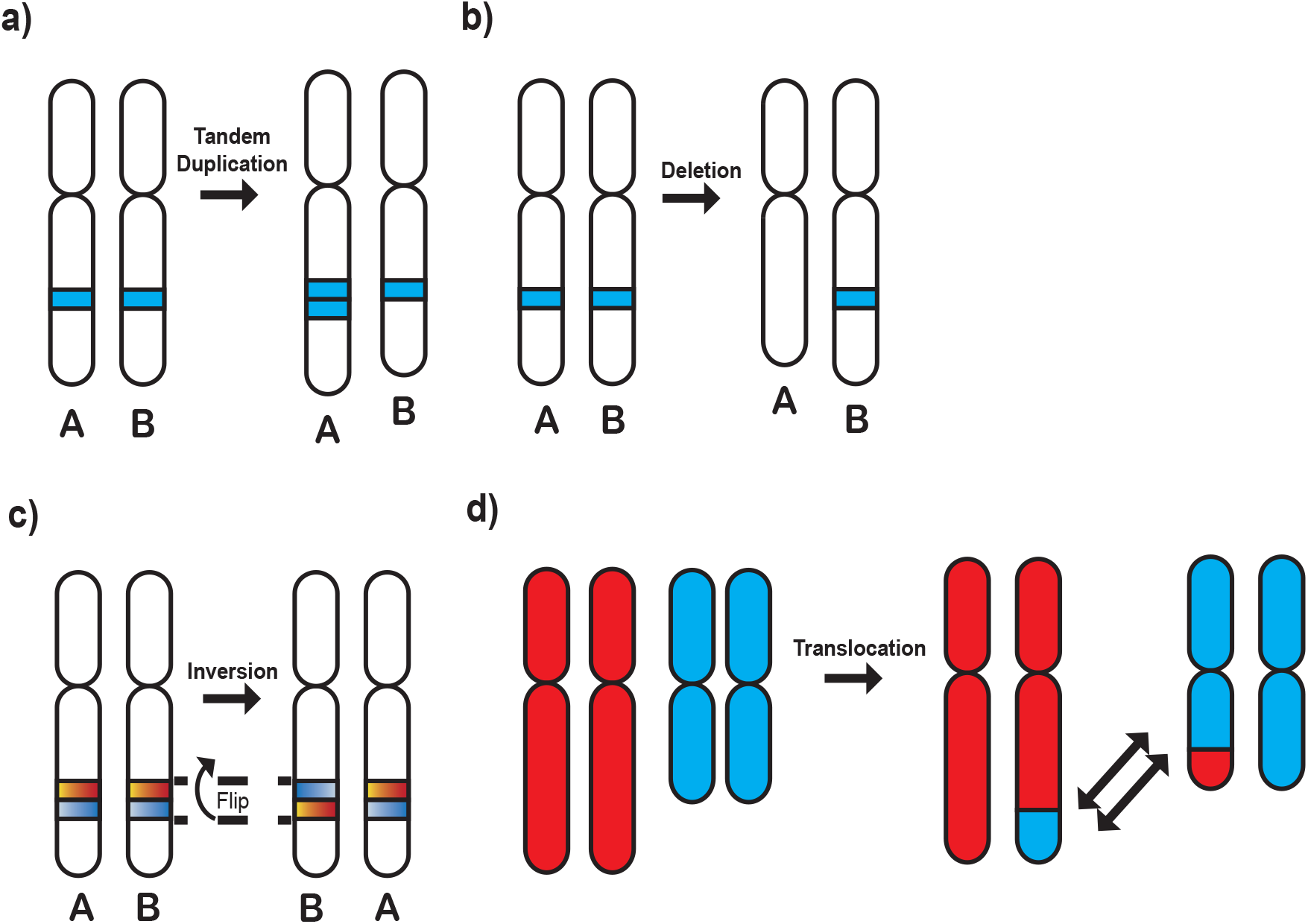
Description of the Structural Variant Classification Schema. Structural variants (SVs) are categorized as tandem-duplications, deletions, inversions, or translocations. **a)** Tandem duplication of a segment containing the A allele. A tandem duplication occurs when a segment is duplicated and inserted adjacent to the original chromosomal segment. **b)** Deletion of the segment containing the A allele. A deletion occurs when there is a loss of genetic material from a chromosome. **c)** An inversion of the segment containing the B allele. An inversion occurs when a segment breaks off and reattaches in a reverse orientation within the same chromosome. **d)** A translocation of a chromosomal segment. A translocation event occurs when a piece of one chromosome breaks off and some (or all) fragments from that piece re-attach to either another chromosome or to a different locus of the same chromosome.

An example of a whole-genome sequenced bone cancer with a highly rearranged genome that contains chromosomes with clustered events as well as chromosomes with only non-clustered events is shown in **Figure 4*a***. For instance, in this sample, chromosome 12 contains a high number of SV breakpoints in close proximity to one another (**Figure 4*b*)** and the SV pattern of this chromosome can be summarized in a vector with 32 components containing a high number of clustered SVs (**Figure 4*d*)**. In contrast, chromosome 8 has SV breakpoints randomly scattered throughout the chromosome (**Figure 4*c*)** and the SV pattern of chromosome 8 is exclusively one of non-clustered SVs (**Figure 4*e***).

**Figure 4.**
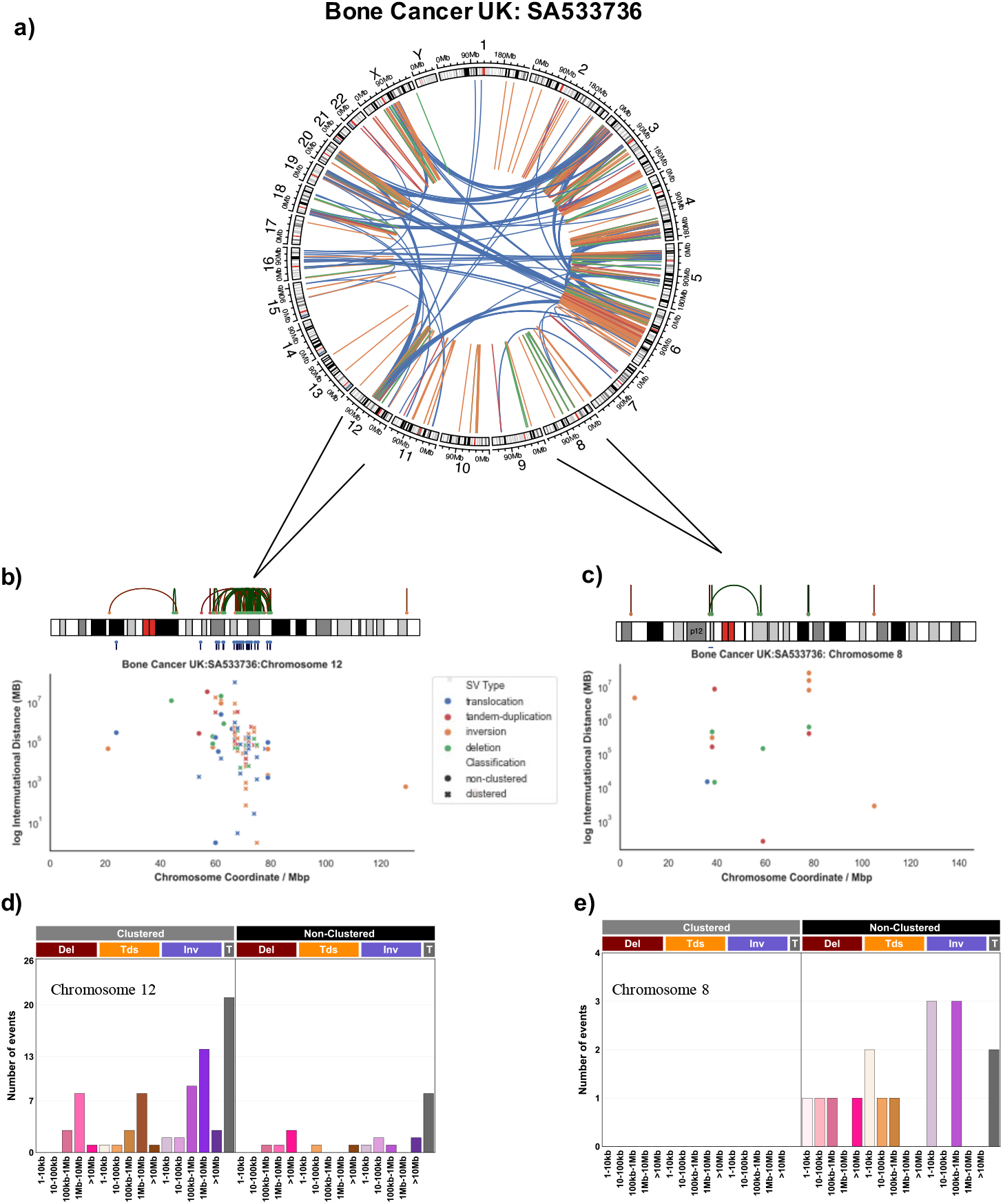
Classifying Structural Variants into Mutational Vectors. a) An example of a bone cancer sample from PCAWG with a highly rearranged genome consisting of both clustered and non-clustered structural variants (SVs) is shown as a Circos plot representation. b) Zooming into SVs specifically found on chromosome 12 in the bone cancer sample. SVs are shown as a linear representation (top) and as a rainfall plot (bottom). The rainfall plot depicts all breakpoints on chromosome 12 according to their genomic coordinate (x-axis) and the log_10_ inter-mutational distance (y-axis), which is the distance to the breakpoint immediately preceding it. The tendency of breakpoints to cluster in a specific genomic region on chromosome 12 due to a chromothripsis event is evident in all representations. c) Zooming into SVs specifically found on chromosome 8 in the bone cancer sample. SVs are shown as a linear representation (top) and as a rainfall plot (bottom). The rainfall plot depicts all breakpoints on chromosome 8 according to their genomic coordinate (x-axis) and the log_10_ inter-mutational distance (y-axis), which is the distance to the breakpoint immediately preceding it. There are no clustered SVs on chromosome 8 as, per the SV classification schema, clustering requires a minimum of 10 breakpoints in a segment of a chromosome. d) The SV classification schema is applied to the SVs found on chromosome 12 in the bone cancer sample. SVs are classified by the event type (denoted by color) and are binned according to the size of the event (0 – 10kb, 10kb – 100kb, 100kb – 1Mb, 1Mb – 10Mb, and >10Mb). e) The SV classification schema is applied to the SVs found on chromosome 8 in the bone cancer sample. SVs are classified by the event type (denoted by color) and are binned according to the size of the event (0 – 10kb, 10kb – 100kb, 100kb – 1Mb, 1Mb – 10Mb, and >10Mb).

### Input Data for Classifying Structural Variants

SigProfilerMatrixGenerator allows examining SV data that contains genomics information for each of the two breakpoints of a structural variant. In principle, the tool can process files in browser extensible data paired-end (BEDPE) format that, at a minimum, contain the following six columns: *chrom1, start1, end1, chrom2, start2*, and *end2*. Here, the genomics coordinates of the first breakpoint are annotated as *chrom1, start1*, and *end1*, while the genomics coordinates of the second breakpoint are provided as *chrom2, start2*, and *end2*. If the type of SV has been predetermined, then its annotation can be provided using a column named *svclass*. Otherwise, the columns *strand1* and *strand2*, which indicate the strands of the read mate-pairs, are required. If the mates are on the same chromosome, the convention followed is inversion (+/- or -/+), deletion (+/+), and tandem-duplication (-/-). If mates are on different chromosomes, the SV is automatically classified as a translocation. SigProfilerMatrixGenerator supports SV in BEDPE format, which is utilized by most bioinformatics tools for detecting SVs, as well as being the native output files from BRASS [21].

## DISCUSSION

The newly developed version of SigProfilerMatrixGenerator allows transforming a set of mutational catalogues of copy-number changes and structural rearrangements into matrices amenable to decomposition, including, subsequent mutational signature analysis. The tool provides support for two previously developed [7, 21] classification schemas for large mutational events. Further, the tool also delivers an extensive plotting functionality that seamlessly integrates with matrix generation to visualize the majority of output in a single analysis. SigProfilerMatrixGenerator is the first tool to provide support for the 48 channel CNV schema across a wide variety of popular tools for detecting CNV. Importantly, this schema can be applied across several data modalities, including whole-genome sequencing, whole-exome sequencing, RNA-sequencing, single-cell sequencing approaches, and genotyping microarrays. In addition, SigProfilerMatrixGenerator is the first Python package that provides support for the 32 channel SV schema in a fast and intuitive manner with minimal preprocessing.

## CONCLUSION

A breadth of computational tools exists for exploring the patterns for small mutational events, including our initial implementation of SigProfilerMatrixGenerator [8]. However, to the best of our knowledge, there are currently no tool for exploration and visualization of large mutational events. We recently demonstrated that a classification of CNVs into 48 channels provides the means to better elucidate and understand the mutational processes operative in human cancer [7]. Similarly, we and others have previously demonstrated that the classification of SVs into 32 channels can be used to understand the mutational processes giving rise to SVs across multiple cancer types [19]. Our newly developed version of SigProfilerMatrixGenerator provides the capability to examine these classification schemas from cancer genomics sequencing data. The tool can scale to large datasets and will serve as foundation to future analysis of both mutational patterns and mutational signatures of large mutational events.

## AVAILABILITY AND REQUIREMENTS

**Project name:** SigProfilerMatrixGenerator

**Project home page:** https://github.com/ Al exandrovLab/SigProfil erMatrixGenerator, https://github.com/AlexandrovLab/SigProfilerMatrixGeneratorR

**Operating system(s):** Unix, Linux, and Windows

**Programming language:** Python 3 and R

**Other requirements:** None

**License:** BSD 2-Clause “Simplified” License

**Any restrictions to use by non-academics:** None

## ABBREVIATIONS

BEDPE: browser extensible data paired-end
CNV: copy number variation
DBS: doublet base substitution
ecDNA: extrachromosomal DNA
ID: small insertions and deletions
LOH: loss of heterozygosity
SBS: single base substitution
SV: structural variation
TCN: total copy-number
WGS: whole-genome sequencing

## DECLARATIONS

### Ethics approval and consent to participate

Not applicable.

### Consent for publication

Not applicable.

### Availability of data and materials

Data sharing is not applicable to this article as no datasets were generated or analyzed during the current study.

### Competing interests

LBA is a compensated consultant and has equity interest in io9, LLC. His spouse is an employee of Biotheranostics, Inc. LBA is also an inventor of a US Patent 10,776,718 for source identification by non-negative matrix factorization. LBA declares U.S. provisional applications with serial numbers: 63/289,601; 63/269,033; 63/366,392; 63/367,846; 63/412,835. All other authors declare that they have no competing interests.

### Funding

This work was supported by the US National Institute of Health grants R01ES030993-01A1, R01ES032547-01, and R01CA269919-01 to LBA as well as Cancer Research UK Grand Challenge Award C98/A24032. This work was also supported a Packard Fellowship for Science and Engineering. The funders had no roles in study design, data collection and analysis, decision to publish, or preparation of the manuscript.

### Authors’ contributions

AK developed the Python and R code with assistance from RV, MB, AA, ENB, and MDG. AA, CDS, and NP tested and evaluated the performance of the code. CDS, NP, and LBA developed the copy number classifications schema. AK wrote the manuscript with assistance from RV, MB, CDS, AA, and MDG. LBA supervised the overall development of the code and writing of the manuscript. All authors read and approved the final manuscript.

## Acknowledgements

The computational development reported in this manuscript have utilized the Triton Shared Computing Cluster at the San Diego Supercomputer Center of UC San Diego.

